# Photovoltaic Implant Simulator Reveals the Resolution Limits in Subretinal Prosthesis

**DOI:** 10.1101/2022.06.30.498210

**Authors:** Zhijie Charles Chen, Bing-Yi Wang, Anna Kochnev Goldstein, Emma Butt, Keith Mathieson, Daniel Palanker

**Affiliations:** Department of Electrical Engineering, Stanford University, Stanford, CA, USA; Department of Physics, Stanford University, Stanford, CA, USA; Institute of Photonics, Dept. of Physics, University of Strathclyde, Glasgow, Scotland, UK; Department of Ophthalmology, Stanford University, Stanford, CA, USA; Hansen Experimental Physics Laboratory, Stanford University, Stanford, CA, USA

## Abstract

**Objective:** PRIMA, the photovoltaic subretinal prosthesis, restores central vision in patients blinded by atrophic age-related macular degeneration (AMD), with a resolution closely matching the 100 µm pixel size of the implant. Improvement in resolution requires smaller pixels, but the resultant electric field may not provide sufficient stimulation strength in the inner nuclear layer (INL) or may lead to excessive crosstalk between neighboring electrodes, giving low contrast stimulation patterns. We study approaches to shaping the electric field in the retina for prosthetic vision with higher resolution and improved contrast.

**Approach:** We present a new computational framework, RPSim, that efficiently computes the electric field in the retina generated by a photovoltaic implant with thousands of electrodes. Leveraging the PRIMA clinical results as a benchmark, we use RPSim to predict the stimulus strength and contrast of the electric field in the retina with various pixel designs and stimulation patterns.

**Main results:** We demonstrate that by utilizing monopolar pixels as both anodes and cathodes to suppress crosstalk, most patients may achieve resolution no worse than 48 µm. Closer proximity between the electrodes and the INL, achieved with pillar electrodes, enhances the stimulus strength and contrast and may enable 24 µm resolution with 20 µm pixels, at least in some patients.

**Significance:** A resolution of 24 µm on the retina corresponds to a visual acuity of 20/100, which is over 4 times higher than the current best prosthetic acuity of 20/438, promising a significant improvement of central vision for many AMD patients.

## 1 Introduction

Recent years has seen the development of retinal prostheses to treat retinal degenerative diseases, including the atrophic age-related macular degeneration (AMD) and retinitis pigmentosa (RP), for which there is no other cure. The first clinical trial of the subretinal implant PRIMA (Pixium Vision, Paris, France) with 100 µm pixels demonstrated the highest prosthetic acuity of 20/438, corresponding to 1.04 times the pixel pitch. The average visual acuity was 20/490 (20/438 to 20/564 range), corresponding to 1.17±0.13 of the pixel size [1]. The close match of the prosthetic acuity to the fundamental sampling limit set by the pixel size indicates that smaller pixels may provide higher resolution. Prosthetic visual acuity can also be limited by electrical crosstalk between the neighboring pixels. In the PRIMA system, such crosstalk is suppressed by the return electrodes located around the active electrode in each pixel, connected in a hexagonal mesh (Figure 1A).

**Figure 1:**
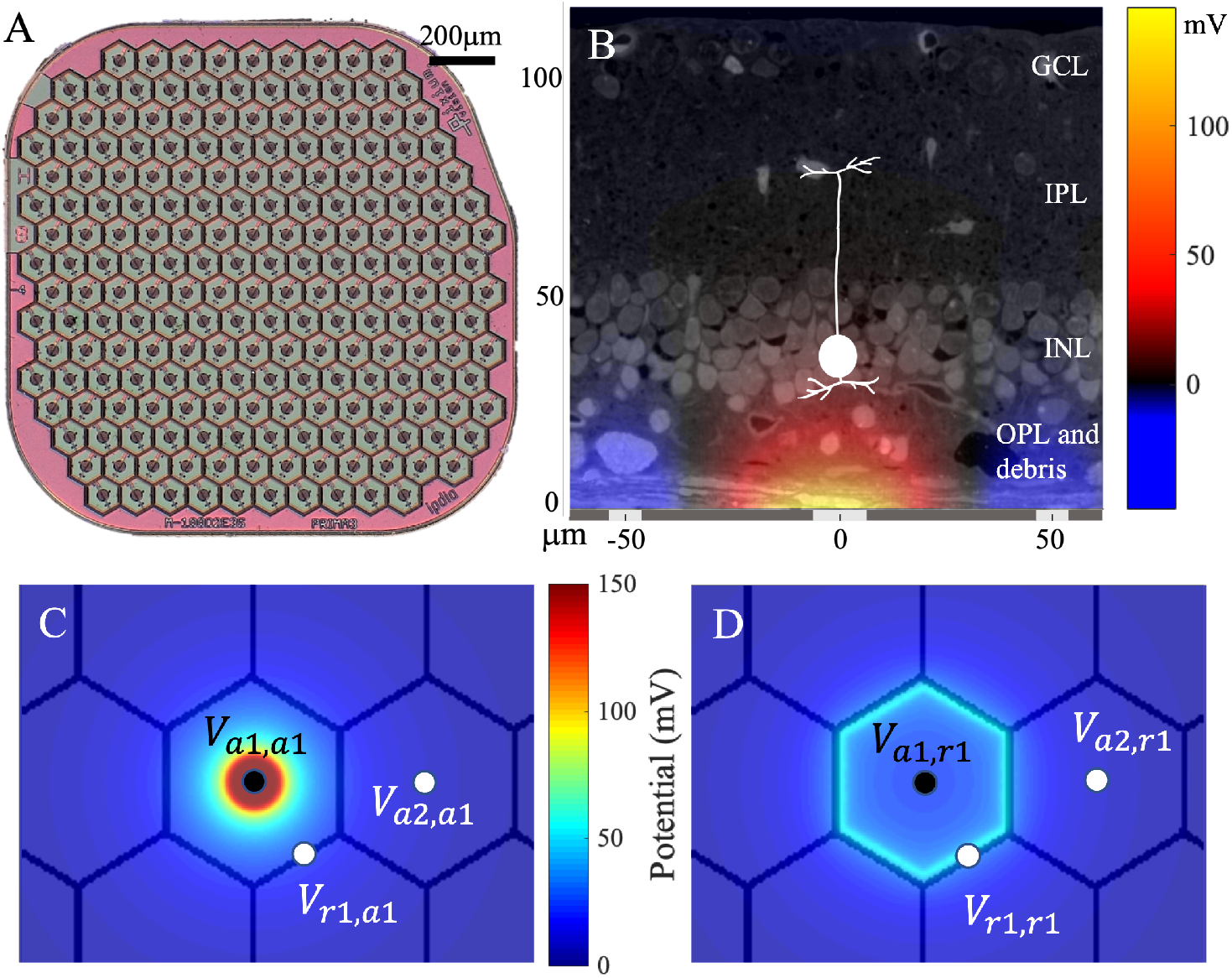
(A) The PRIMA subretinal implant with 100 µm photovoltaic bipolar pixels. (B) Electric potential generated by the 100 µm pixel injecting 1 µA of current, overlaid with a histological representation of the retina with a debris layer and a bipolar cell highlighted. (C, D) Elementary fields of (C) the active and (D) the return electrode of a bipolar pixel.

To stimulate the retina, the active electrode injects electric current into the tissue to generate a voltage drop between the dendritic end of the bipolar cell in the inner nuclear layer (INL) and its axonal arbor in the inner plexiform layer (IPL, Figure 1B). Depolarization of the axonal terminals leads to opening of the voltage-sensitive calcium channels in the axonal terminals, which increases the release rate of glutamate into the retinal ganglion cell synapses [2]. With bipolar pixels, the penetration depth of the electric field (E-field) in tissue is constrained by the distance between the active and the return electrodes. With bipolar pixels smaller than 40 µm, retinal stimulation, even in rats, requires prohibitively high current, exceeding the charge injection limit of one of the best electrode materials - sputtered iridium oxide films (SIROF) [3]. In human patients, where an additional debris layer of about 35 µm in thickness separates the INL from the implant [4], the stimulation threshold is even higher. Multiple designs have been proposed to increase the penetration depth of electric field into the retina, including pillars [5–7], honeycomb wells [3], and current steering with monopolar pixels for E-field shaping [8–10].

Studies of such designs involve modeling the dynamic E-field in the retina, which should include simulation of the circuit dynamics in each pixel, combined with calculation of the E-field in electrolyte for a specific current configuration. With a few pixels, it can be accomplished using the finite element method (FEM). However, such an approach is computationally intractable with a large number of electrodes and multiple levels of current injection at each electrode.

Here we present the Retinal Prosthesis Simulator (RPSim), a framework capable of efficient computation of the dynamic E-field in the retina generated by an implant with thousands of photovoltaic pixels coupled via a common electrolyte. We use RPSim to relate the clinical results with the PRIMA system to the electrical stimulation threshold and contrast performance in the presence of a 35 µm debris layer. With the PRIMA clinical data as a benchmark, we then model the performance of subretinal implants with 40 µm and 20 µm monopolar pixels, where current steering via transient returns (Figure 6A) and the pillar electrodes (Figure 6B) are utilized to further improve the contrast and the stimulation strength. The modeling results show the expected limits of resolution and contrast with a photovoltaic subretinal prosthesis. This computational framework can also be adapted to model other retinal prosthetic systems, or a general neural stimulation with large multi-electrode arrays (MEAs).

## 2 Methods

### 2.1 E-Field Modeling from Elementary Fields

Volume conduction of electric current in biological tissues is described by Poisson’s equation, the linearity of which enables synthesizing the E-field by a basis of elementary fields, each of which is generated by one electrode injecting a unitary current individually. The resulting E-field in the medium can be computed efficiently by superposing all the elementary fields weighted by the currents at their corresponding electrodes [10]. In the monopolar configuration, the current is injected from an active electrode and sinks into the common global return. In the bipolar configuration, each segment of the return mesh also corresponds to an elementary field to account for the non-uniform current distribution across the return mesh (Figure 1C and 1D), where the nominal ground is set at infinity. Note that by the Kirchhoff’s Current Law, after superposing the elementary fields according to the circuit dynamics, the net current to the infinity is always zero. All elementary fields are computed with FEM in COMSOL Multiphysics 5.6 (COMSOL, Inc., Burlington, MA) using the AC/DC module.

To further improve the computational efficiency, we exploit the spatial invariance of the elementary fields in the bipolar configuration, based on the observation that the elementary fields are laterally translated versions of each other. Therefore, we only need to compute two elementary fields with FEM – one for the active electrode and one for the return – instead 410 fields (2 for each of the 205 pixels in Figure 1A). In addition, the elementary field for the active electrode is axially symmetric, making the computation, data storage, and synthesis of the elementary fields very efficient. The elementary fields above the pillars are found to be isomorphic to those with flat pixels (*r* = 0.9936), subject to a scaling difference of 0.92. For convenience of exploring various pillar heights, we use the elementary fields of the flat pixels shifted up by the height and scaled by 0.92, as a proxy for the elementary fields of pillars.

### 2.2 Circuit Model of the Implant

To compute the circuit dynamics with up to thousands of interrelated pixels in the implant, RPSim uses Xyce, a SPICE-compatible circuit simulator designed for extremely large circuit problems, as its kernel solver [11]. RPSim provides users with an interface in Python to configure the circuit, and calls Xyce through the PySpice module [12]. Our code is opensource on Github [13].

As illustrated in Figure 2, RPSim comprises four major components: the electrolyte, the electrode-electrolyte interface, the pixel and the photocurrent driver. The coupling of pixels in an array through the electrolyte is quantified by the resistance matrix **R** and the multidimensional Ohm’s Law ***V*** = **R*I***, where ***V*** and ***I*** are vectors consisting of the potential and current at all electrodes, respectively [14]. RPSim converts the resistance matrix to an equivalent mesh of resistors to comply with available features in Xyce, as described in the Appendix. The electrode-electrolyte interface is modeled by a surface capacitance of 6 mF cm^−2^ based on our previous characterization of SIROF [15]. The circuit consists of two diodes and a shunt resistor of 720 kΩ for the PRIMA bipolar 100 µm pixels, and one diode with 750 kΩ or 1.75 MΩ shunt resistor for monopolar 40 µm and 20 µm pixels, respectively. The photocurrent driver, which is a current pulse generator with an output proportional to irradiance on each pixel, is in parallel with the diodes, and together they mimic the current injection of the photodiodes under illumination [16]. Each component can be independently substituted, and for non-photovoltaic implants, the pixel and the photocurrent driver can be replaced by circuits corresponding to the power mechanism (amplifiers, for example).

**Figure 2:**
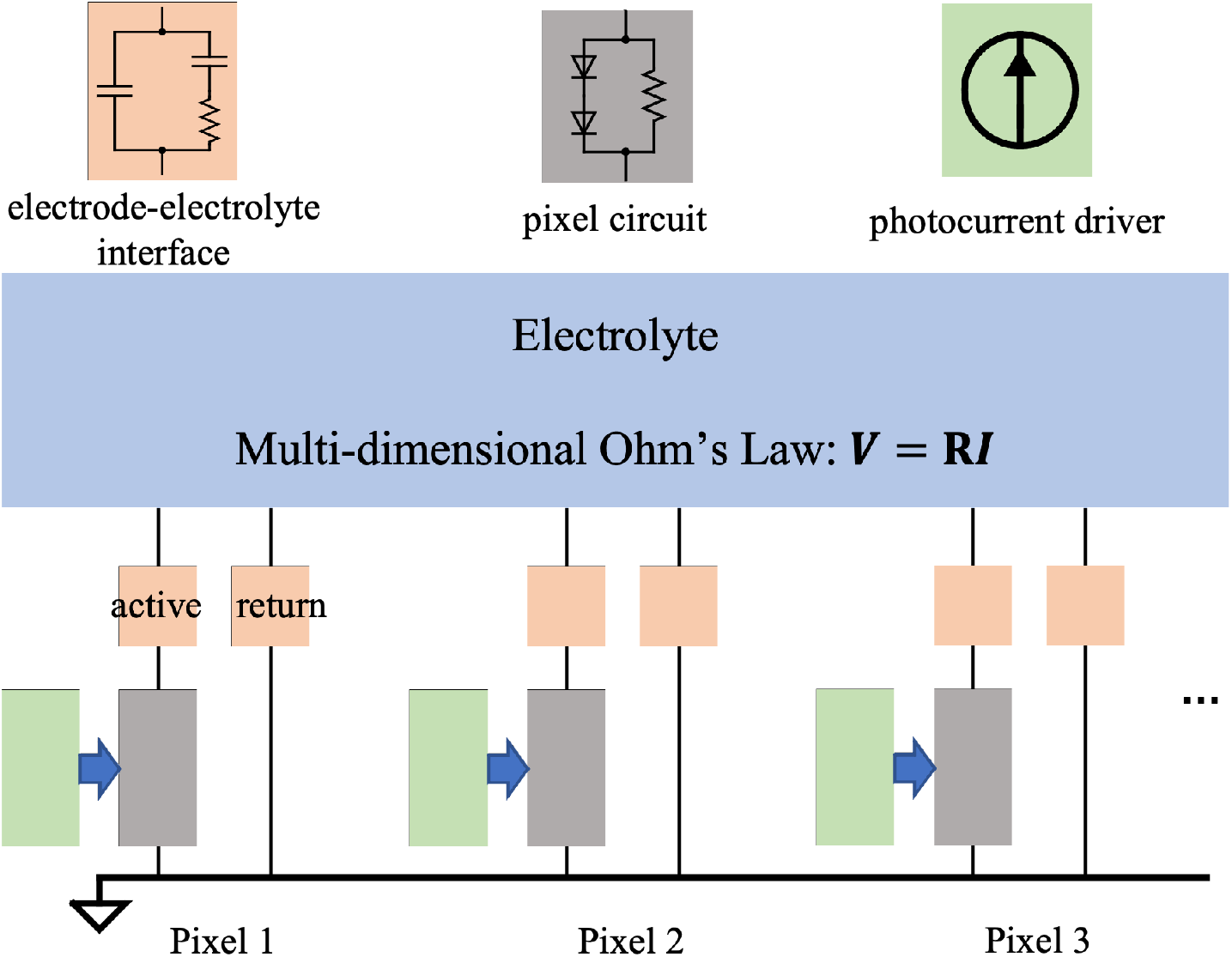
Diagram of the RPSim framework for modeling the dynamic E-field in the retina generated by an implant with up to thousands of photovoltaic pixels. The sub-circuit in each block can be adjusted independently. Current generated by an illuminated photovoltaic pixel is represented by a current source (green) in parallel with the pixel circuit in the dark (gray).

### 2.3 In-Vitro Validation of the Model

The modeling framework has been validated previously for the monopolar pixels array [14]. To validate the bipolar configuration, we compared the RPSim modeling result with in-vitro measurement and with an FEM model in COMSOL. For the measurement, we placed a PRIMA implant with 100 µm pixels in a Petri dish filled to a depth of 4 mm with NaCl solution of 1.52 mS cm^−1^ conductivity. A micro-pipette, with an opening of approximately 5 µm, was placed 17 µm above the center pixel (Figure 3A). The 880 nm laser beam was projected at 3 mW mm^−2^ irradiance with 9.6ms pulses repeated at 30Hz, over the whole array (full field) or a spot covering about half of the pixels (Figure 3B). The potential waveform shown in Figure 3C was averaged over 100 measurements to reduce the noise. For the FEM model, we constructed the geometry of the in-vitro measurement in COMSOL and used the electrochemistry module and the electric circuits interface from the AC/DC module to calculate the voltage waveform with the same circuit parameters as RPSim. Unlike the discretized return electrodes in RPSim, where a uniform current density is assumed in each segment, the COMSOL model treats the return mesh as a continuous surface, where current distribution is governed by the dynamics of the electrode-electrolyte interface.

**Figure 3:**
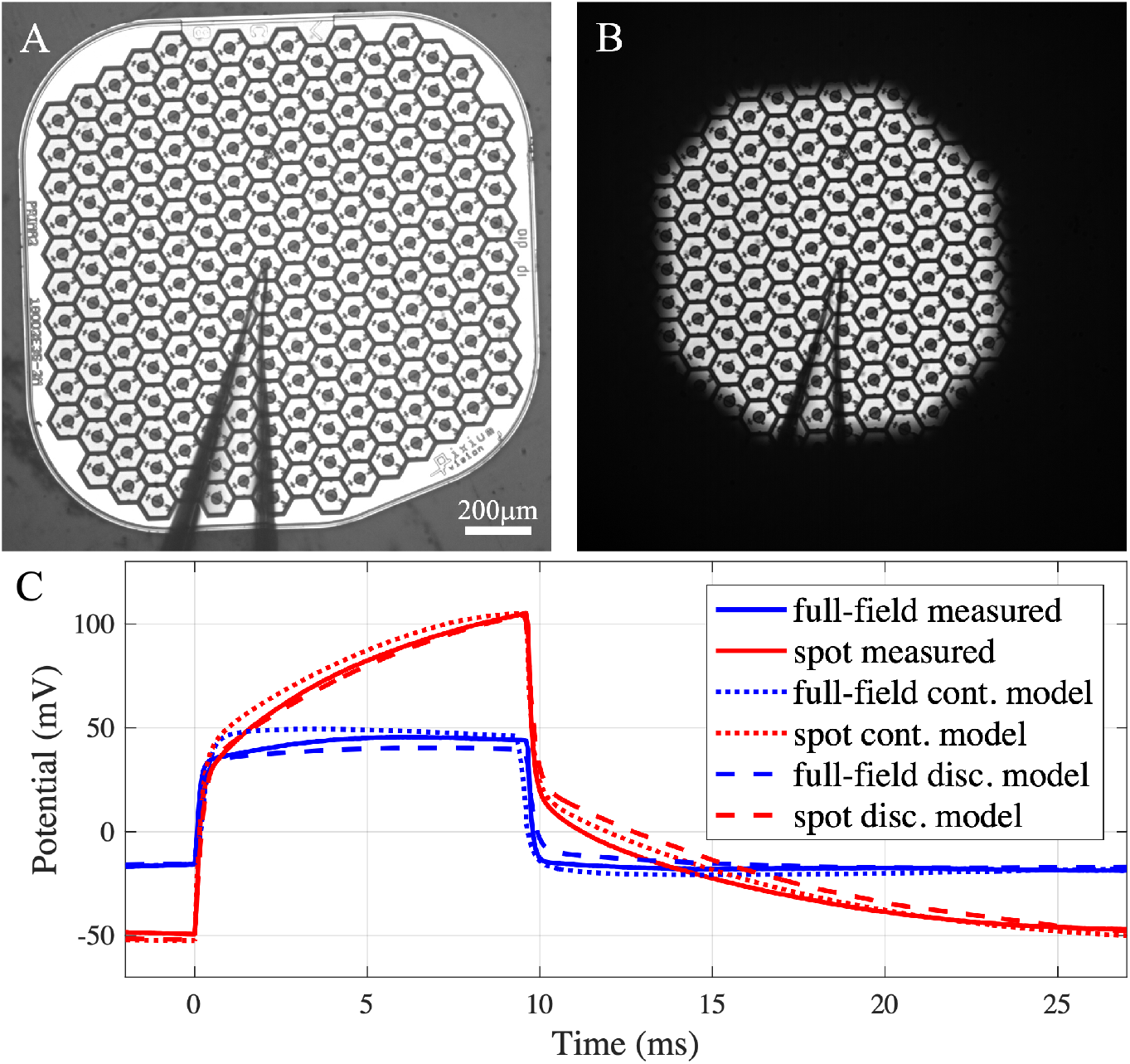
In-vitro validation of the model. (A,B) A PRIMA implant tested in (A) fullfield and (B) spot illumination, both at 3 mW mm^−2^, using 9.6 ms pulses at 30 Hz repetition rate. The outline of the micro-pipette, placed above the center pixel, can be seen. (C) The potential waveform measured 17 µm above the center pixel under the two illumination patterns (solid lines). Modeling results by (dotted lines) FEM in COMSOL, where the return mesh is a continuous domain, and (dash lines) by RPSim, where the return mesh is discretized into pixels.

### 2.4 Modeling E-Field in the Retina

We model the degenerate retina (including the INL, IPL and GCL, as shown in Figure 1B) as a 100 µm-thick layer with a conductivity of 1 mS cm^−1^ [17]. In RCS rats, the BC bodies are separated from the implant by only about 5 µm [18], while in humans, the INL is separated from electrodes by a debris layer of about 35 µm in thickness. We assume all the retinal layers have the same conductivity. A large volume representing the vitreous humor is placed above the retina, with a conductivity of 15 mS cm^−1^ [19, 20]. The voltage drop between the somatic and the axonal ends of a BC governs its neural response [2], and we assume that the axonal terminals are located in the middle of IPL, at 87 µm above the implant surface (Figure 1B). The potential at the somatic end of a BC is defined by the average potential across its cross section, which is assumed to be 10 µm in diameter. To account for the probabilistic BC layout, in which the potential maximum may be anywhere relative to the cell, we estimate the stimulation strength of an image by convolving the potential map with a 20 µm-diameter “electrical receptive field” and finding the maximum after convolution. We also assume that the BCs located closest to the implant will be activated first since they experience the highest potential. Their cell bodies are located in the INL, but the dendrites might reside below - in the OPL, as discussed in Section 3.2.

Following the PRIMA clinical protocol, we model the stimulation threshold by projecting a 1.6 mm spot on the implant at 10 Hz repetition rate, with pulse width and irradiance drawn from the clinical strength-duration (S-D) curves shown in Figure 4A [4]. RPSim calculates the current injection of each pixel as a function of time (Figure 4B), and the voltage drop across the BCs is computed by superimposing the elementary fields (Figure 4C). We average the voltage drop over the pulse width and the electrical receptive field to find the maximum stimulation strength of an image, which we define as the threshold. Then, we model the electric field for an acuity test using the Landolt C with 120 µm gap width (1.2 × pixel size) projected at 3 mW mm^−2^ irradiance with 9.8 ms pulses repeated at 30 Hz. The voltage contrast is defined by (1 − *V*_gap_) */* (1 − *V*_stroke_), where *V*_gap_ is the average voltage drop in the middle of the gap, and *V*_stroke_ - over the semicircle opposite to the gap. Next, the PRIMA implant is replaced by a monopolar array with 821 40 µm pixels or 2806 20 µm pixels. Each type was modelled with and without pillars of 30 µm height. The pillar height is designed to bring the electrodes sufficiently close to the BCs (as close as in RCS rats) through the debris layer. The Landolt C test is repeated with these pixels, using the gap width of 1.2 times the pixel sizes (48 µm and 24 µm) and the same irradiance and pulse duration. The stimulation strength (voltage drop across BCs) and contrast are compared to those with the PRIMA implant.

**Figure 4:**
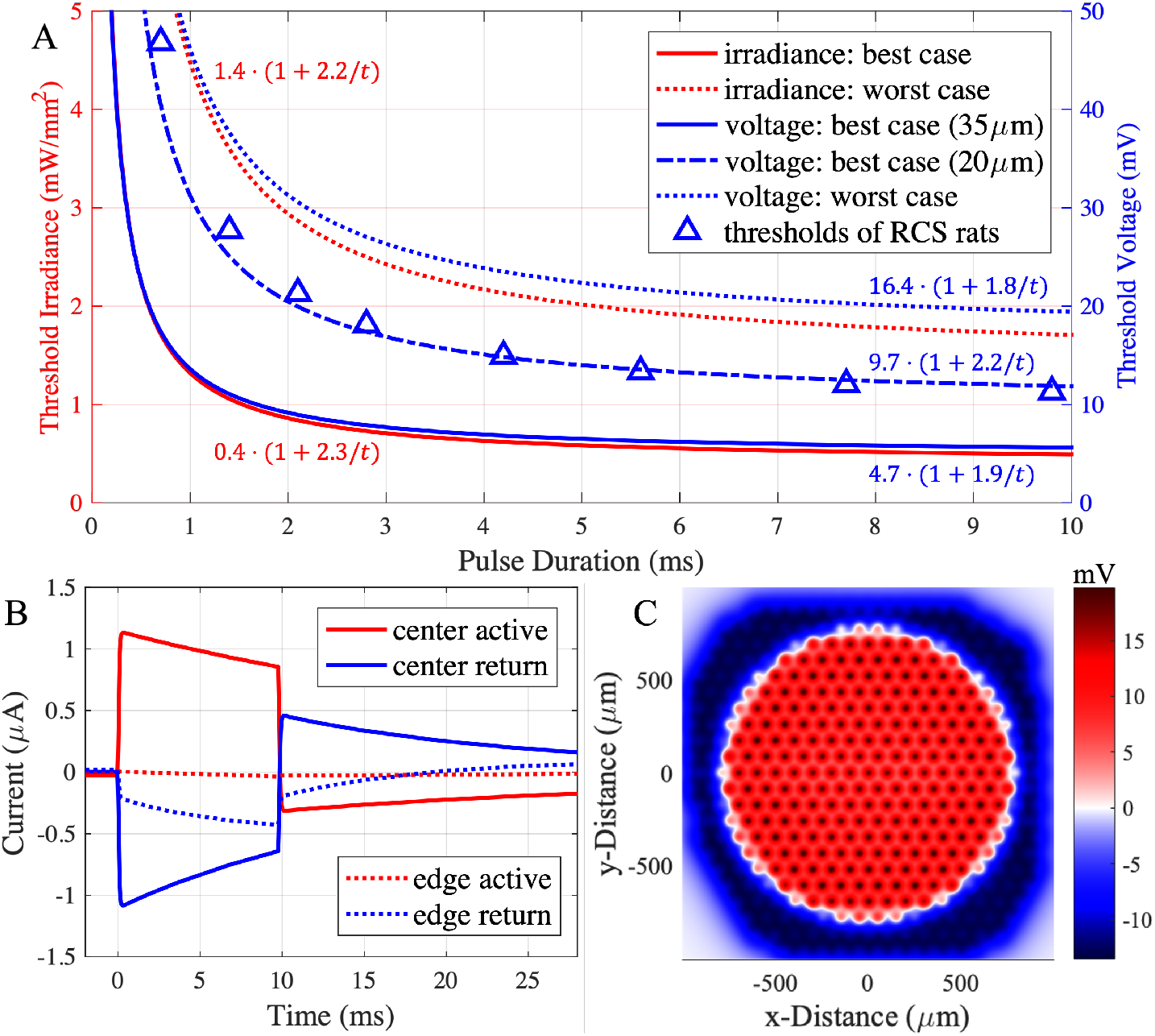
(A) The S-D curves of the best-case (lowest) and the worst-case (highest) irradiance thresholds in the first clinical trial. The voltage S-D curves are computed for the bestcase at the electrophysiologically realistic position of the dentrites at 20 µm, or the “naive” one at 35 µm. For the worst case, voltage threshold is calculated for 35 µm separation of the INL. Thresholds of RCS rats are shown for comparison [22]. (B) Current injection waveforms of two pixels under the 1.6 mm illumination spot in the PRIMA clinical threshold test, calculated by RPSim for the worst-case with 9.8 ms pulse duration at 1.71 mW mm^−2^ irradiance. (C) Potential across the BCs corresponding to the stimulus in (B).

## 3 Results

### 3.1 In-Vitro Validation of the Model

In Figure 3C, the voltage waveform measured under full-field illumination resembles the square pulse of light, and the amplitude is well matched by both (RPSim and COMSOL) models. Under the spot illumination, however, the potential increases during the pulse, which is the result of transition from the equipotential boundary condition at the beginning of the pulse to a uniform current density on the return mesh [21]. At the onset of the pulse, current from the active electrodes in illuminated pixels follows the path of least resistance and sinks to the nearest return segments. As charge accumulates and voltage on the return electrode rises in the illuminated region, current is driven to the more remote segments in the unilluminated area, leading to deeper penetration of the E-field into the electrolyte and hence higher potential, as measured via the pipette. The COMSOL FEM computes the dynamics of the continuous current distribution on the return mesh, which accurately captures the characteristic ramp up, but takes 12 min to simulate each pulse cycle of 33 ms on a desktop computer (Intel Core i7-8700K, 3.7 GHz, 64 GB memory). In contrast, RPSim divides the return mesh into hexagonal segments of single pixels, assuming a uniform current density in each segment, and the circuit solver determines the current distribution among the segments. Such discrete approximation achieves similar agreement with the measurement, yet the simulation takes less than 10 s per pulse cycle, improving the computational efficiency by almost two orders of magnitude.

### 3.2 Stimulation Threshold and Contrast with the PRIMA Implants

Clinical trials with the PRIMA implants demonstrated variability in the stimulation thresholds among patients by a factor of about 3.5 [4]. Figure 4A shows the lowest and the highest strength-duration (S-D) curves among them (rheobase of 0.4 and 1.4 mW mm^−2^), which we use as the best- and the worst-case benchmarks, respectively, to estimate the range of responses with the future implants. We convert the two S-D curves from irradiance into voltage across bipolar cells by first computing the current injection of each pixel with RPSim (Figure 4B) and then finding the maximum voltage drop across the BCs above the implant (Figure 4C). The resulting voltage S-D curves of the best- and the worst-case patients yield rheobases of 4.7 mV and 16.4 mV, respectively, assuming the BCs extend from 35 µm to 87 µm above the implant.

The rheobase in the worst-case patient is about twice the rodent rheobase [22], which may result from reduced neural excitability in the badly degenerate retina. However, the best-case rheobase, 4.7 mV, appears to be too low to stimulate the BCs. Literature indicates that the BC threshold is closer to 10 mV [17, 23, 24], which is consistent with the dynamics of the L-type Ca^2+^ channel [2], the dominant voltage-gated ion channel at the axonal terminal [25]. Such discrepancy may be reconciled by assuming the BC dendrites extend below the INL into the debris layer, thereby reducing the distance to the implant and leading to a higher voltage drop. If we assume a stimulation threshold at about 10 mV for the BCs, the dendrites should be separated from the implant by about 20 µm. This refined S-D curve in Figure 4A yields a rheobase of 9.7 mV. This curve matches very well the thresholds in rats [22], shown by triangles in Figure 4B.

Another figure of merit to consider is the resolution of the Landolt C font used in the clinical trials for assessment of the visual acuity. Patients with the PRIMA 100 µm pixels, on average, were able to resolve the Landolt C with a 120 µm gap (1.2 × pixel size) [1]. The corresponding projection pattern is shown in Figure 5A, and the voltage drop across BCs for the starting height of 35 µm and 20 µm - in Figure 5C and 5D, respectively. The stimulation strengths are found to be 31 mV and 62 mV for the two cases, exceeding the stimulation threshold of even for the worst case (19.7 mV). The contours in Figure 5C and 5D outline the corresponding stimulation thresholds for 10 ms pulses in the best case, using the “naive” (5.7 mV) and the “electrophysiologically realistic” (11.7 mV) assumptions for 10 ms pulses, respectively. The two resulting contours are very similar, demonstrating that the model predictions regarding visibility of the letter do not depend strongly on this assumption. To illustrate the contrast of the letter, we plot the BC voltage drop averaged over the stroke of the C, as a function of the polar angle *θ* (Figure 5B), where *θ* = 0 corresponds to the gap. The contrast is shown by the ratio between the average over the semicircle opposite to the gap (*θ* ∈ [0.5*π, π*]) and the minimum of the plot inside the gap (*θ* = 0). The worst case yields a contrast of 94 %, and the best case – 100 %, highlighting the well-confined electric field with 100 µm bipolar pixels.

**Figure 5:**
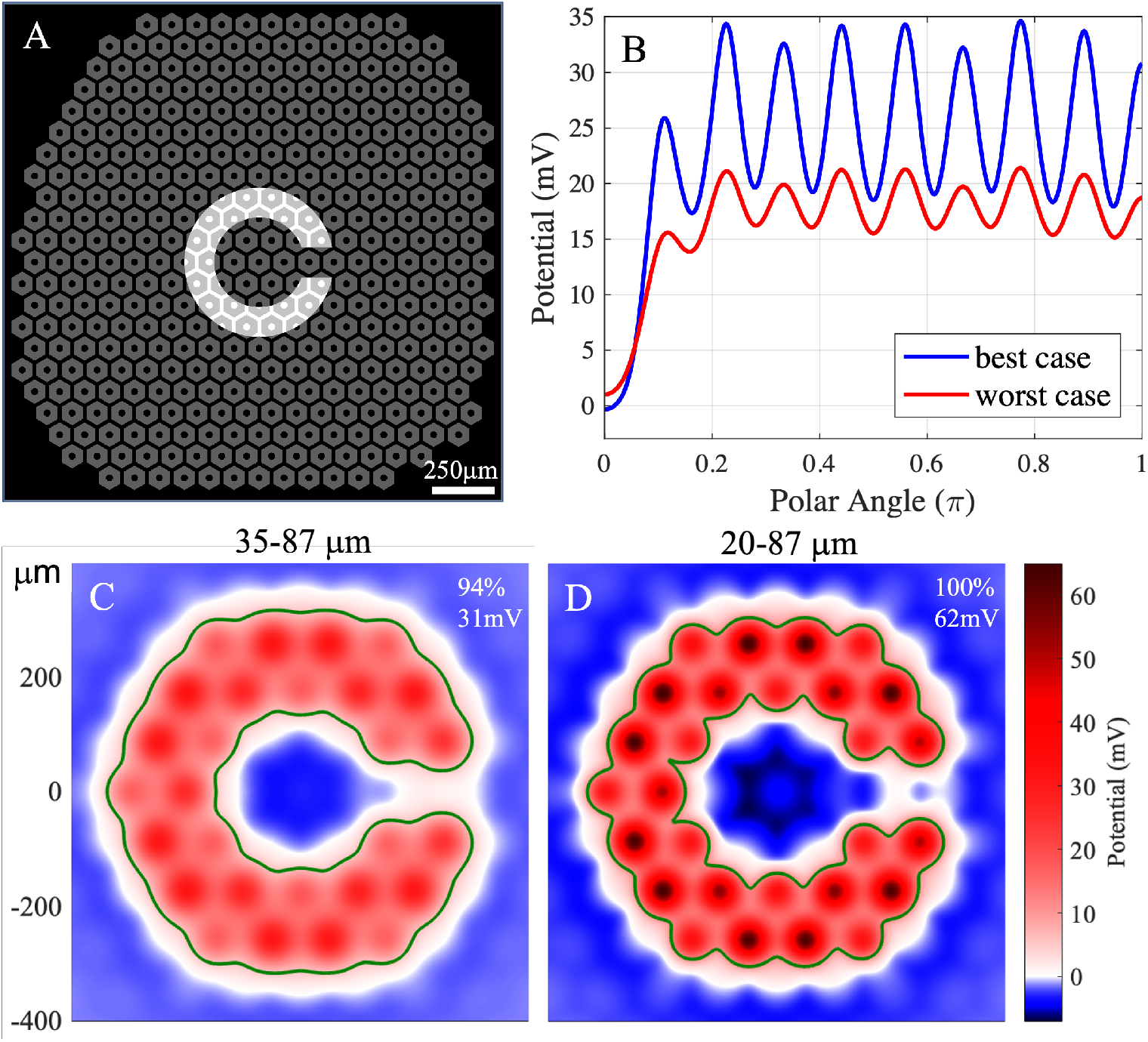
(A) The 100 µm PRIMA array overlaid with the average minimal resolvable Landolt C, having a 120 µm-wide gap (1.2 × pixel size) illuminated by 3 mW mm^−2^, 9.8 ms pulses at 30 Hz repetition rate. (B) Potential averaged across the stroke width of the C, as a function of the polar angle about the center, where the gap is at 0 degree. (C, D) Potential across the BCs for the worst (C) and the best (D) cases. Labels in the top-right corner show the contrast and the stimulation strength of each figure. The contours outline the area above the stimulation threshold in the best case, calculated by the “naive” conversion (35 µm height, 5.7 mV) in (C) and the electrophysiologically realistic one (20 µm height, 11.7 mV) in (D).

### 3.3 Stimulation Strategies with Small Pixels

Flat bipolar pixels smaller than 40 µm cannot effectively stimulate the BCs even in RCS rats, where there is no debris layer [3], and even less so in the human retina. Therefore, a different electrode configuration should be used to enable on one hand, deep enough penetration of electric field into the INL, while on the other hand, sufficient field confinement to avoid excessive interference (crosstalk) from the neighboring pixels, thereby preserving high contrast of the stimulation pattern. Here, we model the performance of the 40 µm and 20 µm monopolar pixels with two strategies – current steering (Figure 6A), and pillar electrodes (Figure 6B).

**Figure 6:**
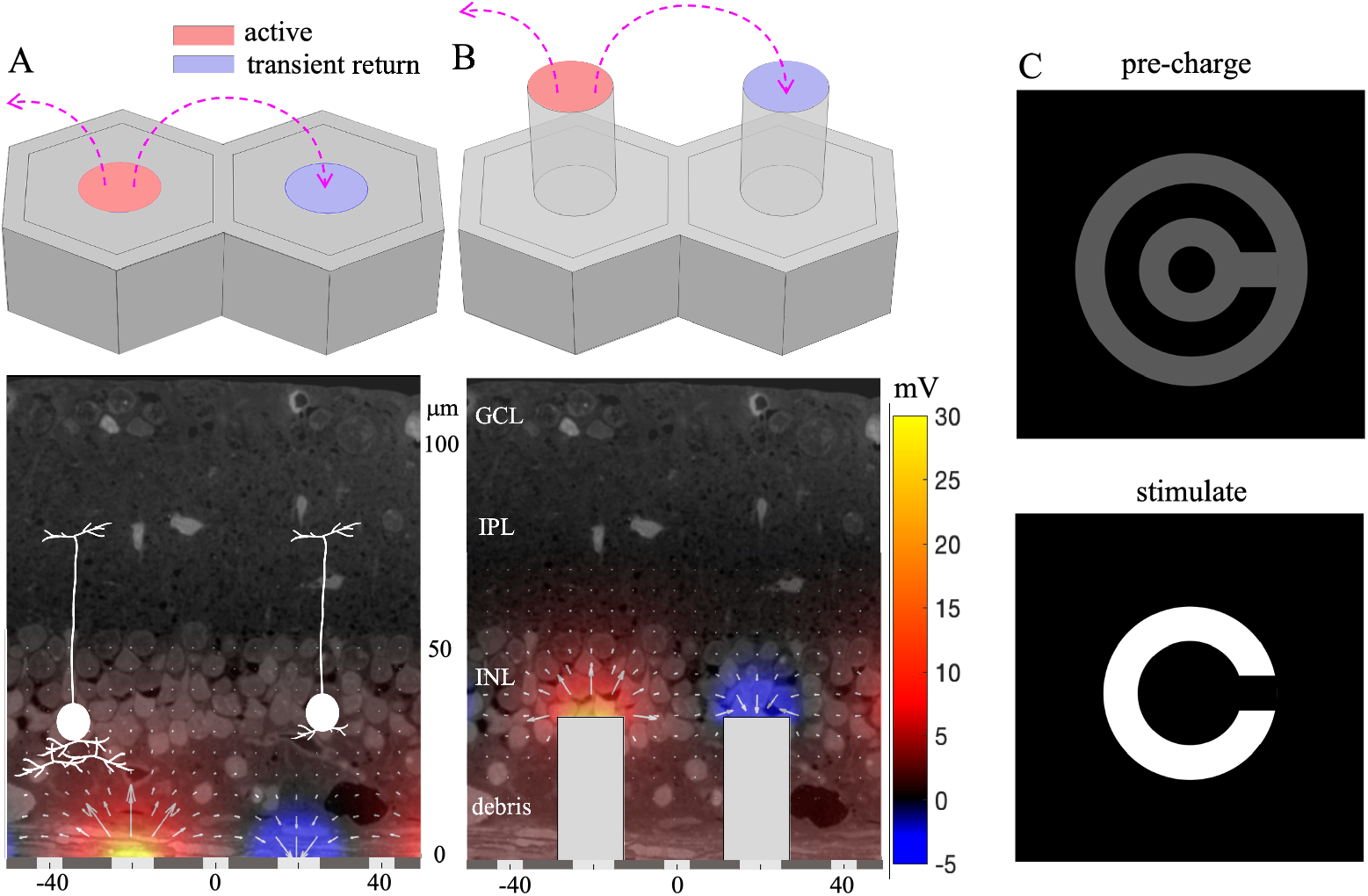
(A) Schematic of the 40 µm flat monopolar pixels (top) and the electric field they generate, overlaid with a histological micrograph of the retina and two bipolar cell diagrams illustrating the best and the worst cases (bottom). (B) 40 µm monopolar pixels with pillars penetrating through the debris layer. (C) Pre-charging the dark pixels around the illumination pattern with 35 % of the stimulation irradiance (top) to enhance their conductance during the stimulation phase (bottom).

The current steering strategy with monopolar pixels is based on the use of the active electrodes as both the anodes and cathodes, i.e. current is injected into electrolyte from the illuminated pixels and collected via the dark pixels. Conductance of the dark pixels is provided by two mechanisms: through the shunt resistor in parallel with the photodiode (Figure 2), and through the diodes under forward voltage bias. For the first mechanism, the shunt resistor should be conductive enough to sink the cathodal current, but not too conductive so the photocurrent is primarily injected into the electrolyte rather than shorted via the shunt during stimulation. The shunt resistor also speeds-up a discharge of the electrodes between the pulses, thereby increasing the charge injection at higher repetition rates [26]. For 0.7 ms and 9.8 ms pulses at 30 Hz, the shunt resistors that maximize the charge injection were found to be about 10 and 3 times the access resistance of a single pixel, respectively, although the maximum charge injection is not very sensitive to the shunt value. Since long pulses are more voltage-limited by the photodiodes, and a smaller shunt resistance is more favorable for the field confinement, the optimal shunt was determined to be about 3 times the access resistance - 750 kΩ and 1.75 MΩ for 40 µm and 20 µm pixels, respectively.

Conduction through photodiodes can be enhanced by pre-charging the designated pixels prior to stimulation. Charge accumulation at the capacitive electrode-electrolyte interface creates a forward bias across the diode, which increases its conductance and hence the return current in the stimulation phase. Stronger return current on the neighboring pixels enhances the field confinement and the associated contrast of the electrical pattern [14]. Pre-charging should be performed below the stimulation threshold, so that by itself it will not elicit neural responses. With 30 Hz repetition rate and a 9.8 ms stimulus pulse, 23.5 ms is available within each stimulation period for pre-charging. For confinement of electric field in a pattern, a rim of pixels around the stimulation pattern can be exposed to light at a sub-threshold irradiance level between the stimulation pulses, so they become forward-biased prior to the stimulation phase. Such a pre-charging pattern can be produced by widening the stimulation pattern by 1 pixel and subtracting the stimulation pattern, as shown in Figure 6C. During the stimulation pulse, these pixels are not exposed to light, so they serve as enhanced transient returns for the illuminated pixels, which emit the photocurrent.

Another approach to enhance penetration of electric field into the retina is to use pillars that penetrate through the debris layer and bring the electrodes closer to the INL, due to cellular migration [27]. In modeling this approach, we assume a 5 µm distance between the electrodes and BCs, as observed in RCS rats [18]. The two strategies (current steering and pillars) can be combined.

### 3.4 Stimulation Strength and Contrast with Monopolar Pixels

We model the voltage drop across BC with 40 µm and 20 µm monopolar pixels in a Landolt C pattern, where the gap is set at 48 µm and 24 µm, respectively, corresponding to the achievable resolution of 1.2 × pixel size in PRIMA patients [1]. Figure 7 shows the potential maps with 40 µm flat pixels for the worst and the best cases, and of pillars, with and without the pre-charged transient returns. The contrast and the stimulation strength are shown in the top-right corner of each map. With flat pixels and without optically-enhanced current steering, both cases exhibit contrast below 50 %. Pre-charging the edge pixels improves the contrast to a level comparable to that with PRIMA 100 µm pixels, at the price of reduced stimulation strength. As a result, the stimulation strength for the worst case, 26 mV, is close to the stimulation threshold of 19.7 mV from the clinical data. With pillars, the stimulus is much stronger and the contrast is high (75 %), even without the enhanced current steering - with optical pre-charging it reaches 100 %.

**Figure 7:**
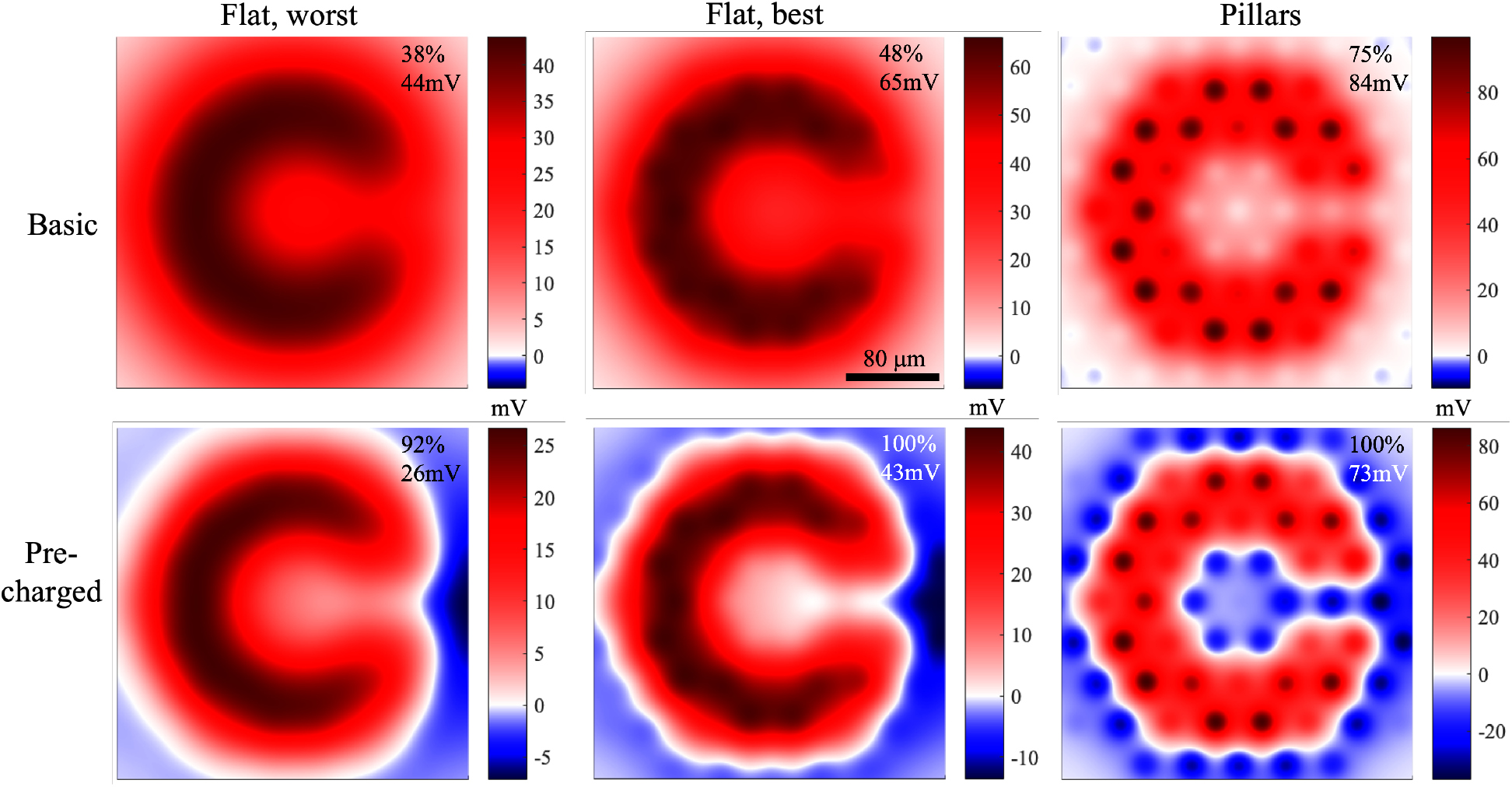
Voltage drop across the BCs generated by the 40 µm monopolar pixels, with the Landolt C of 48 µm gap projected by 3 mW mm^−2^, 9.8 ms pulses at 30 Hz repetition rate. Results are shown for the worst and the best cases with flat pixels, and for pillars, with and without the pre-charging.

A similar set of field maps is shown in Figure 8 for 20 µm pixels and the corresponding Landolt C size. In the worst case, the stimulation strength with flat pixels is below the threshold, even without the optical current steering. In the best case, only pre-charging provides a satisfactory contrast, but the stimulation strength, 15 mV, is close to the threshold of 11.7 mV. With pillars and without current steering, the contrast is mediocre (68 %), but it can reach 100 % with the current steering, although the stimulation is rather weak (24 mV). Table 1 summaries the contrast and stimulation strength of all the configurations mentioned above, while the gap of the Landolt C is always 1.2 times the pixel size. As expected, pillars yield stronger stimulation and better contrast than flat pixels under the same conditions.

**Table 1:** Contrast and amplitude (voltage/threshold) generated by 100 µm, 40 µm and 20 µm pixels for the smallest resolvable C (gap width = 1.2 × pixel size) under 3 mW mm^−2^ irradiance, with 9.8 ms pulses at 30 Hz repetition rate. Stimulation thresholds are 11.7 mV and 19.7 mV in the best and the worst cases, respectively.

| | | 100 $\mu\text{m}$ PRIMA | | 40 $\mu\text{m}$ pixels | | 20 $\mu\text{m}$ pixels | |
| --- | --- | --- | --- | --- | --- | --- | --- |
|  |  | contrast (%) | voltage/threshold | contrast (%) | voltage/threshold | contrast (%) | voltage/threshold |
| Flat, worst | basic | 94 | 1.6 | 38 | 2.2 | 36 | 0.83 |
|  | pre-charged |  |  | 92 | 1.3 | 75 | 0.45 |
| Flat, best | basic | 100 | 5.3 | 48 | 5.6 | 44 | 2.2 |
|  | pre-charged |  |  | 100 | 3.7 | 87 | 1.3 |
| Pillars, worst | basic |  |  | 75 | 4.3 | 68 | 1.6 |
|  | pre-charged |  |  | 100 | 3.6 | 100 | 1.2 |
| Pillars, best | basic |  |  | 75 | 7.9 | 68 | 3.0 |
|  | pre-charged |  |  | 100 | 6.8 | 100 | 2.2 |

**Figure 8:**
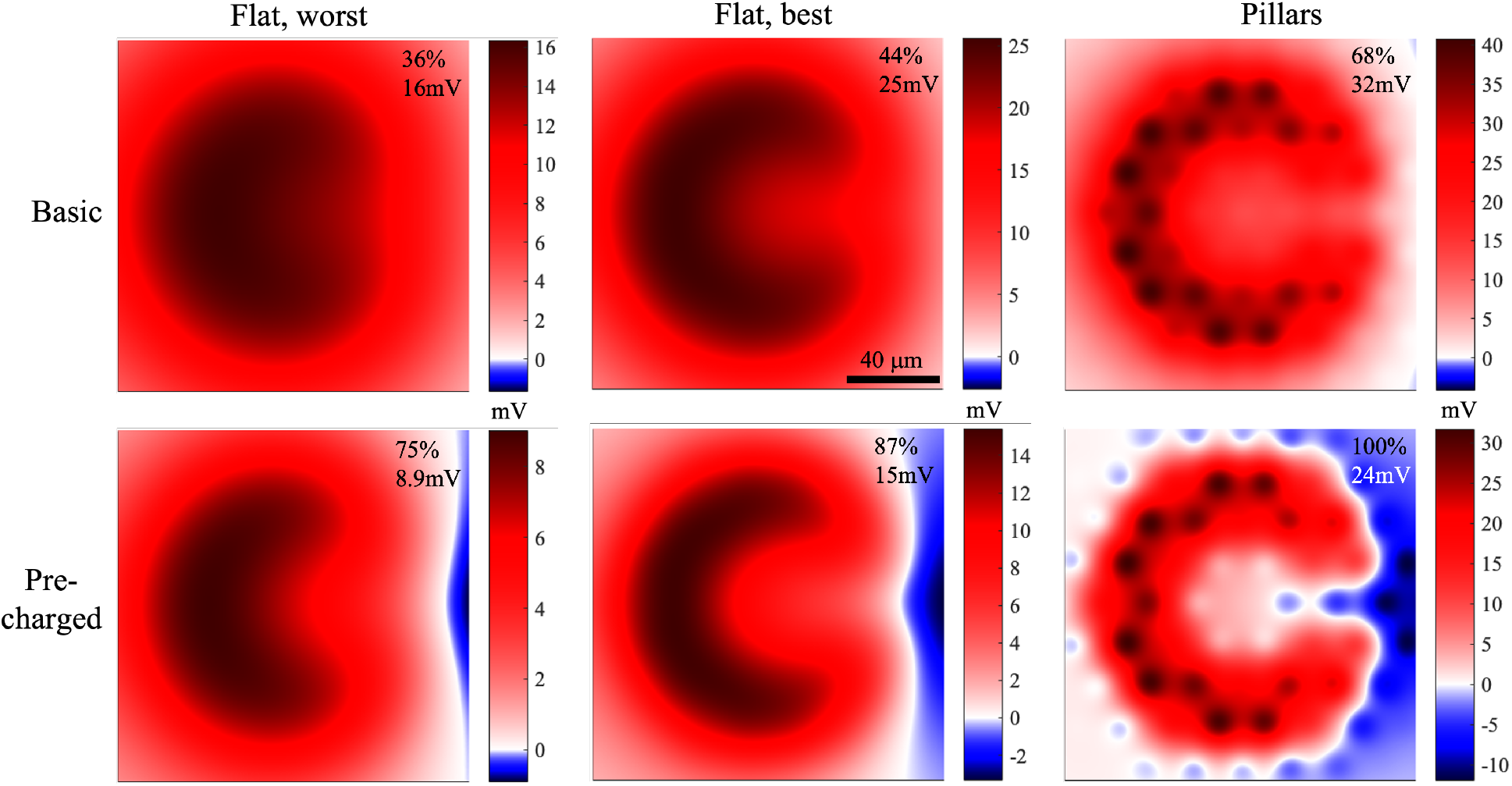
Voltage drop across the BCs generated by the 20 µm monopolar pixels, with the Landolt C of 24 µm gap projected by 3 mW mm^−2^, 9.8 ms pulses at 30 Hz repetition rate. Results are shown for the worst and the best cases with flat pixels, and for pillars, with and without the pre-charging.

### 3.5 20 µm Pixels with Optimal Shunt as a Universal Hardware

According to Table 1, the 1.2 × pixel size resolution with 40 µm pixels may be achieved in all patients, while achieving 24 µm resolution with 20 µm pixels is difficult due to rather low amplitude and contrast even with pillars, which maybe sufficient only for the best case. However, the question remains what is the best implant configuration for the 40 µm resolution? Figure 9 shows the optimal configurations for resolving 48 µm gap of Landolt C in the worst-case patient. With 40 µm pixels, sufficient contrast is achieved only with pre-charging the edge pixels. With 20 µm pixels, however, similar stimulation strength and contrast levels are achieved with only the basic current steering through the shunt resistors. The 20 µm pixel array provides a better contrast than the 40 µm pixels, under the basic current steering, due to the shorter distance between the illuminated pixels injecting the current and the unilluminated neighbors, which serve as returns. Pillars enhance the stimulus strength to a similar extent in both cases. Avoiding the need for pre-charging greatly simplifies the device operation. All configurations yield sufficient stimulation strength and contrast even for the worst case, as demonstrated by the contours for the worst-case stimulation threshold of 19.7 mV. Contrast with flat 20 µm pixels and no pre-charging (72 %) is a little lower than with 40 µm and optical current steering (92 %). The remedy, however, can be a weaker pre-charging to improve the contrast without sacrificing too much in stimulation strength. Therefore, implants with 20 µm monopolar pixels and pillars may serve as a universal hardware for all patients, which can support resolution of no worse than 48 µm, and in some patients may be as good as 24 µm, corresponding to the visual acuity of 20/100.

**Figure 9:**
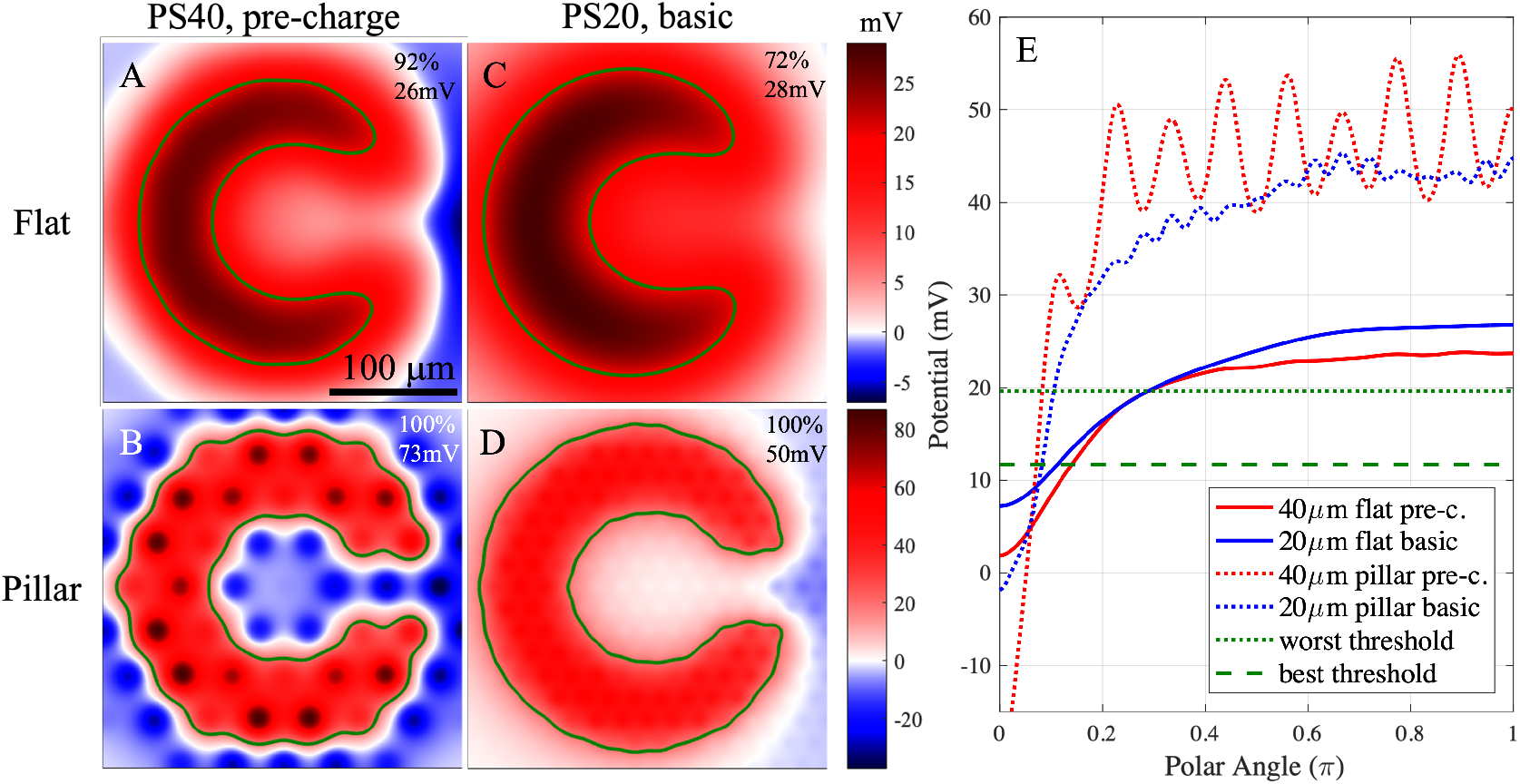
(A-D) Voltage drop across the BCs generated by the Landolt C pattern of 48 µm gap at 3 mW mm^−2^, 9.8 ms pulses at 30 Hz repetition rate. (A) Flat 40 µm pixels with pre-charging in the worst case; (B) 40 µm pillars with pre-charging; (C) flat 20 µm pixels without pre-charging; (D) 20 µm pillars without pre-charging. (E) Potential in (A-D) averaged across the stroke width of the C as a function of the polar angle about the center, where the gap is at 0 degree. Horizontal dash and dotted lines show the stimulation thresholds for the best and the worst cases, respectively.

## 4 Discussion

The RPSim tool we developed is a highly efficient approach to modeling the electric field in electrolyte generated by a large-scale MEA, at the core of which is the superposition of the elementary fields. Although the FEM is still required to generate the elementary fields, they only need to be computed once, and any E-field in the medium can be obtained by simple scaling and summation of them. Furthermore, if only some linear metrics of the electric field are of interest, for example, the voltage between the two ends of the BCs in our case, we can store just these metrics in each elementary field, instead of the potential in the entire 3-dimensional (3-D) volume, which further improves the computational efficiency. The only premise of this framework is the linear volume conduction of electric current in the medium, which is valid for most biological tissues even those with anisotropic conductivity [28–30], and is not affected by nonlinearity of the electrode dynamics or other parts of the MEA circuitry. RPSim is modularized by design and hence can be easily adapted for modeling other retinal prosthetic systems or other neural modulators.

The boundary condition for the current on a capacitive electrode-electrolyte interface transitions from equipotential at the beginning of the pulse to a current density proportional to the capacitance per unit area in the steady state [21]. With a homogeneous electrode surface, this means a uniform current density. Such a transition can be described as a family of eigenmodes of charge and current distribution, each decaying with its own time constant. The eigenmodes with stronger spatial variation decay faster due to the less resistive pathway for the charge to equilibrate in the local vicinity. RPSim assumes a uniform current density within the return segment of each pixel, which is less precise than the complete FEM, where the return mesh is modeled as a continuous surface. However, such discretization only results in omitting the fast-decaying eigenmodes, while the current redistribution on the array scale is governed by the slowest ones. Therefore, RPSim was able to achieve a similar accuracy as the FEM, but in less than 2% of the computation time.

To verify whether the voltage drop across BCs determines the retinal stimulation threshold, and hence our assumption that the best-case patient’s dendrites start in the debris layer, we measured the irradiance thresholds in RCS rats with the PRIMA implants of 100 µm and 75 µm pixels and converted the irradiances into voltages across BCs. Despite the 2.5-fold difference in the irradiance thresholds with these two pixel sizes, the voltage drop across BCs was very similar [22], supporting the view that they indeed determine the stimulation threshold. More details regarding the rodent experiments are described in the companion paper [22]. Despite the difference between the electrophysiologically realistic conversion assuming the dendrites at 20 µm, and the “naive” conversion at 35 µm, their predictions about the irradiance thresholds and visibility contours with 75 µm bipolar pixels were also very similar, because the threshold derived from either assumption applies likewise to predictive conversions under the same assumption. Such similarity indicates that our model is not very sensitive to this paired assumption about the BC location and stimulation threshold, and hence the prediction is robust.

High-resolution neural stimulation requires both stimulation strength and lateral confinement of E-field to suppress the crosstalk between neighboring electrodes. With a flat monopolar array, where all electrodes inject current of the same polarity, field confinement is very limited. On the other end of the spectrum, bipolar pixels with a return electrode inside each pixel over-confine the field and may not reach the stimulation threshold for neurons located further away from the implant than the distance between the active and the return electrodes. Current steering in monopolar arrays provides a configurable middle ground between the two extremes, depending on the amount of current sunk to the transient returns. The basic current steering through the shunts in our system provides a baseline confinement, which is sufficient in some cases (Figure 9C and 9D, for example), while providing a simple operation of the device. Contrast can be further enhanced by a software-controlled pre-charging, if stronger field confinement is desired for higher resolution. However, development of a universal pre-charging algorithm for gray-scale images requires further research, and may be affected by the eye movements between the pre-charging and the stimulation phases. One approach to shaping a gray-scale E-field in the retina is by the real-time optimization-based current steering [10].

Nevertheless, current steering alone does not solve the fundamental issue between stimulation strength and field confinement for remote cells, because the lateral confinement also limits the vertical span of the E-fields in the retina. Analogously, current steering trades stimulation strength for more contrast, which would not work if the stimulation is not strong enough. The fundamental solution is 3-D electrodes, such as the pillars demonstrated in this work that bring the electrodes closer to the target cells, and the honeycombs that decouple the field penetration depth from the pixel size while maximizing the stimulation efficiency by vertical alignment of electric field [3]. On the other hand, 3-D electrodes require more complex fabrication and the issues of the retinal integration with the 3-D implant require further study.

## 5 Conclusions

RPSim enables efficient modeling of electric field in the retina generated by thousands of photovoltaic pixels. Using the clinical stimulation thresholds with 100 µm bipolar pixels and the corresponding Landolt C acuity test as a benchmark, such modeling can predict the stimulus strength and contrast with various pixel designs and stimulation patterns. A resolution of 48 µm can be achieved by 40 µm flat monopolar pixels using shunt resistors to sink the current via dark pixels for crosstalk suppression. Optically-controlled pre-charging can further enhance the field confinement by increasing the conductance of the diodes in transient returns. Pillars penetrating through the debris layer reduce the distance between the electrodes and the INL, and hence improve both the stimulation strength and contrast. The modeling indicates that a resolution of 24 µm may be achievable with 20 µm pixels and pillar electrodes.

## Acknowledgements

The authors would like to thank Pixium Vision for providing the PRIMA implants used in this study. Studies were supported by the National Institutes of Health (Grants R01-EY-027786 and P30-EY-026877), the Department of Defense (Grant W81XWH-19-1-0738), AFOSR (Grant FA9550-19-1-0402), Wu Tsai Institute of Neurosciences at Stanford, and unrestricted grant from Research to Prevent Blindness. Photovoltaic arrays were fabricated at the Stanford Nano Shared Facilities (SNSF) and Stanford Nanofabrication Facility (SNF), which are supported by the National Science Foundation award ECCS1542152. K.M. was supported by a Royal Academy of Engineering Chair in Emerging Technology, UK. E.B. was partially supported by the Rhona Reid Charitable Trust.

## Appendix

### Converting Resistance Matrix to Resistor Mesh

Electrodes in an array are coupled via the common electrolyte, and their collective current-voltage (I-V) relationship is described by the resistance matrix **R**, of which the entry at the *m*^th^ row and *n*^th^ column, *R*_*m,n*_, is the cross-pixel resistance from electrode *n* to electrode *m*:

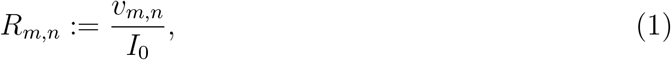

where *v*_*m,n*_ is the average potential on electrode *m* when electrode *n* injects current *I*_0_ individually [14]. However, the resistance matrix is not naturally supported by Xyce or Spice, where the relevant z-parameter network is compatible with neither time-domain analysis nor thousands of ports. Here we show the equivalent conversion from resistance matrix to a mesh of resistors connecting every pair of electrodes.

Let there be *N* electrodes, ***I*** := [*I*_1_, *I*_2_, …, *I*_*N*_]^⊺^ where *I*_*m*_ is the current injection of the *m*^th^ electrode, and ***V*** := [*V*_1_, *V*_2_, …, *V*_*N*_]^⊺^ where *V*_*m*_ is the potential at the *m*^th^ electrode, for all *m* ∈ {1, 2, …, *N*}. By the linearity of volume conduction and Equation (1), we have:

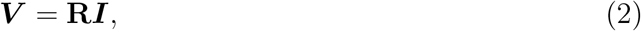

or equivalently,

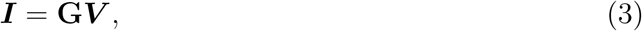

where **G** := **R**^−1^. Let *G*_*m,n*_ be the entry in the *m*^th^ row and *n*^th^ column of **G**.

Instead of the electrolyte, consider a resistor mesh connected between every pair of the electrodes, where *r*_*m,n*_ is the resistor connecting electrode *m* and electrode *n*, and *r*_*m,m*_ is the direct connection between electrode *m* and the ground. Note that such definition requires the symmetry of **R**, which stems from the reciprocity of electromagnetism. The electric current from electrode *m* to electrode *n* through *r*_*m,n*_ is given by

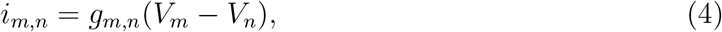

where 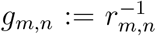. The current directly from electrode *m* to the ground is simply *i*_*m,m*_ = *g*_*m,m*_*V*_*m*_. Therefore, by the linearity of a resistor network, we have:

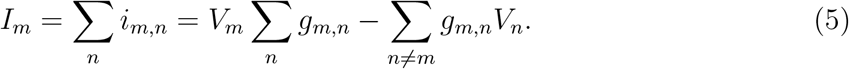

Define

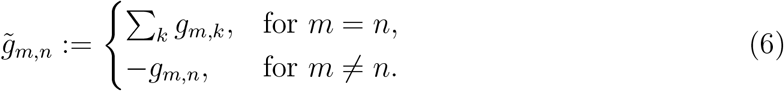

and we have:

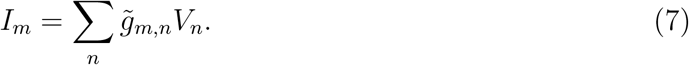

Comparing Equations (3) and (7), the resistor mesh is equivalent to **R**, if

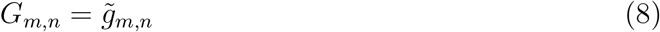

for all *m, n* ∈ {1, 2, …, *N*}. Let us convert the conductance by

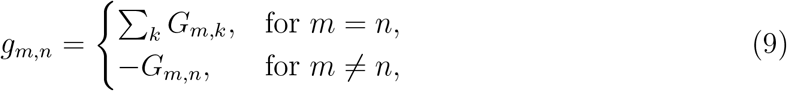

and Equation (8) is satisfied. Equivalently, the conversion can be given in resistance:

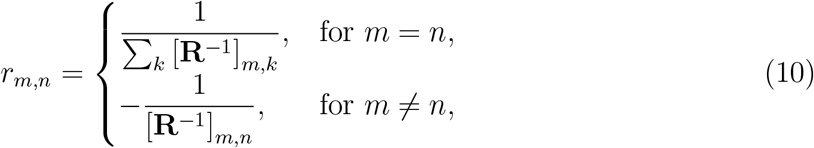

where [**R**^−1^]_*m,n*_ denotes the entry at the *m*^th^ row and *n*^th^ column of **R**^−1^.

## References

[1] Daniel Palanker, Yannick Le Mer, Saddek Mohand-Said, and José-Alain Sahel. Simultaneous perception of prosthetic and natural vision in amd patients. Nature communications, 13(1):1–6, 2022.

[2] Paul Werginz, Bing-Yi Wang, Zhijie Charles Chen, and Daniel Palanker. On optimal coupling of the ‘electronic photoreceptors’ into the degenerate retina. Journal of neural engineering, 17(4):045008, 2020.

[3] Thomas Flores, Tiffany Huang, Mohajeet Bhuckory, Elton Ho, Zhijie Chen, Roopa Dalal, Ludwig Galambos, Theodore Kamins, Keith Mathieson, and Daniel Palanker. Honeycomb-shaped electro-neural interface enables cellular-scale pixels in subretinal prosthesis. Scientific reports, 9(1):1–12, 2019.

[4] Daniel Palanker, Yannick Le Mer, Saddek Mohand-Said, Mahiul Muqit, and Jose A Sahel. Photovoltaic restoration of central vision in atrophic age-related macular degeneration. Ophthalmology, 127(8):1097–1104, 2020.

[5] Eui Tae Kim, Jong-Mo Seo, Se Joon Woo, Jing Ai Zhou, Hum Chung, and Sung June Kim. Fabrication of pillar shaped electrode arrays for artificial retinal implants. Sensors, 8(9):5845–5856, 2008.

[6] A Butterwick, P Huie, BW Jones, RE Marc, M Marmor, and D Palanker. Effect of shape and coating of a subretinal prosthesis on its integration with the retina. Experimental eye research, 88(1):22–29, 2009.

[7] K Koo, S Lee, J Ban, H Jeong, H Park, S Ha, J-M Seo, H Chung, and D Cho. Arrowhead-shaped micro-electrode array on polyimide substrate for retinal prostheses enabling close approach to target cells. In TRANSDUCERS 2009-2009 International Solid-State Sensors, Actuators and Microsystems Conference, pages 342–345. IEEE, 2009.

[8] Thomas C Spencer, James B Fallon, Carla J Abbott, Penny J Allen, Alice Brandli, Chi D Luu, Stephanie B Epp, and Mohit N Shivdasani. Electrical field shaping techniques in a feline model of retinal degeneration. In 2018 40th Annual International Conference of the IEEE Engineering in Medicine and Biology Society (EMBC), pages 1222–1225. IEEE, 2018.

[9] Chung-Yu Wu, Chi-Kuan Tseng, Jung-Hsing Liao, Chuan-Chin Chiao, Fang-Liang Chu, Yueh-Chun Tsai, Jun Ohta, and Toshihiko Noda. Cmos 256-pixel/480-pixel photovoltaic-powered subretinal prosthetic chips with wide image dynamic range and bi/four-directional sharing electrodes and their ex vivo experimental validations with mice. IEEE Transactions on Circuits and Systems I: Regular Papers, 67(10):3273–3283, 2020.

[10] Zhijie Charles Chen, Bing-Yi Wang, and Daniel Palanker. Real-time optimization of the current steering for visual prosthesis. In 2021 10th International IEEE/EMBS Conference on Neural Engineering (NER), pages 592–596. IEEE, 2021.

[11] Eric Keiter, Thomas Russo, Richard Schiek, Heidi Thornquist, Ting Mei, Jason Verley, Karthik Aadithya, and Joshua Schickling. Xyce parallel electronic simulator users’ guide version 7.5. 5 2022.

[12] Fabrice Salvaire. PySpice 1.4.0. https://pyspice.fabrice-salvaire.fr, 2020.

[13] Zhijie Charles Chen, Anna Kochnev Goldstein, and Daniel Palanker. RPsim v1.0.0. https://doi.org/10.5281/zenodo.6774591, June 2022.

[14] Bing-Yi Wang, Zhijie Charles Chen, Mohajeet Bhuckory, Tiffany Huang, Andrew Shin, Valentina Zuckerman, Elton Ho, Ethan Rosenfeld, Ludwig Galambos, Theodore Kamins, et al. Electronic “photoreceptors” enable prosthetic vision with acuity matching the natural resolution in rats. bioRxiv, 2021.

[15] Tiffany W Huang, Theodore I Kamins, Zhijie Charles Chen, Bing-Yi Wang, Mohajeet Bhuckory, Ludwig Galambos, Elton Ho, Tong Ling, Sean Afshar, Andrew Shin, et al. Vertical-junction photodiodes for smaller pixels in retinal prostheses. Journal of neural engineering, 18(3):036015, 2021.

[16] Zhijie Charles Chen, Bing-Yi Wang, and Daniel Palanker. Harmonic-balance circuit analysis for electro-neural interfaces. Journal of Neural Engineering, 17(3):035001, 2020.

[17] P Werginz, H Benav, E Zrenner, and F Rattay. Modeling the response of on and off retinal bipolar cells during electric stimulation. Vision research, 111:170–181, 2015.

[18] Thomas Flores, Georges Goetz, Xin Lei, and Daniel Palanker. Optimization of return electrodes in neurostimulating arrays. Journal of neural engineering, 13(3):036010, 2016.

[19] I Jürgens, J Rosell, and PJ Riu. Electrical impedance tomography of the eye: in vitro measurements of the cornea and the lens. Physiological Measurement, 17(4A):A187, 1996.

[20] Lindenblatt G and Silny J. A model of the electrical volume conductor in the region of the eye in the elf range. Physics in Medicine & Biology, 46(11):3051, 2001.

[21] Zhijie Chen, Lenya Ryzhik, and Daniel Palanker. Current distribution on capacitive electrode-electrolyte interfaces. Physical Review Applied, 13(1):014004, 2020.

[22] Bing-Yi Wang, Zhijie Charles Chen, Mohajeet Bhuckory, Anna Kochnev Goldstein, and Daniel Palanker. Pixel size limit of the prima implants: from humans to rodents and back. bioRxiv, 2022.

[23] Dario A Protti and Isabel Llano. Calcium currents and calcium signaling in rod bipolar cells of rat retinal slices. Journal of Neuroscience, 18(10):3715–3724, 1998.

[24] Heval Benav. Modeling effects of extracellular stimulation on retinal bipolar cells. PhD thesis, Universitätsbibliothek Tübingen, 2012.

[25] Thomas Euler, Silke Haverkamp, Timm Schubert, and Tom Baden. Retinal bipolar cells: elementary building blocks of vision. Nature Reviews Neuroscience, 15(8):507–519, 2014.

[26] David Boinagrov, Xin Lei, Georges Goetz, Theodore I Kamins, Keith Mathieson, Ludwig Galambos, James S Harris, and Daniel Palanker. Photovoltaic pixels for neural stimulation: circuit models and performance. IEEE transactions on biomedical circuits and systems, 10(1):85–97, 2015.

[27] Elton Ho, Xin Lei, Thomas Flores, Henri Lorach, Tiffany Huang, Ludwig Galambos, Theodore Kamins, James Harris, Keith Mathieson, and Daniel Palanker. Characteristics of prosthetic vision in rats with subretinal flat and pillar electrode arrays. Journal of neural engineering, 16(6):066027, 2019.

[28] Socrates Dokos, Gregg J Suaning, and Nigel H Lovell. A bidomain model of epiretinal stimulation. IEEE transactions on neural systems and rehabilitation engineering, 13(2):137–146, 2005.

[29] Mattias Åström, Jean-Jacques Lemaire, and Karin Wårdell. Influence of heterogeneous and anisotropic tissue conductivity on electric field distribution in deep brain stimulation. Medical & biological engineering & computing, 50(1):23–32, 2012.

[30] Syed Salman Shahid, Marom Bikson, Humaira Salman, Peng Wen, and Tony Ahfock. The value and cost of complexity in predictive modelling: role of tissue anisotropic conductivity and fibre tracts in neuromodulation. Journal of neural engineering, 11(3):036002, 2014.

